# Berberine delays onset of collagen induced arthritis through T cell suppression

**DOI:** 10.1101/736264

**Authors:** Alexandra A. Vita, Hend Aljobaily, David O. Lyons, Nicholas A. Pullen

## Abstract

Previous evidence suggests that berberine (BBR), a clinically relevant plant-derived alkaloid, alleviates symptoms of clinically apparent collagen induced arthritis (CIA), and may have a prophylactic role from *in vitro* studies. Thus, we used a CIA model to determine if BBR merits further exploration as a prophylactic treatment for rheumatoid arthritis. Mice were treated with either 1 mg/kg/day of BBR or a vehicle (PBS) control via IP injections from day 0 to day 28, were left untreated (CIA control), or were in a non-arthritic control group. Incidence of arthritis in BBR mice was 40%, compared to 90% in the CIA and 80% in the PBS controls. Populations of B cells and T cells from the spleens and draining lymph nodes were examined from mice across treatment groups on day 14 and from the remaining mice on day 28 when arthritic signs and symptoms were expected to be apparent. BBR-treated mice had significantly reduced populations of CD4^+^ T cells, CXCR5^+^ T_fh_cells, and an increased proportion of T_reg_ at both day 14 and day 28 endpoints, as well as decreased CD28^+^ and CD154^+^ CD4^+^ T cells at day 14. BBR-treated mice also experienced a significant reduction of CD19^+^ B cells in LNs at day 28. Additionally, BBR treatment resulted in significantly lower anti-collagen type II-specific (anti-CII) IgG2a and anti-CII total IgG serum concentrations. These results indicate a potential role for BBR as a prophylactic supplement, and that its effect may be mediated through T cell suppression, which indirectly affects B cell activity.

## I. Intro

Rheumatoid arthritis (RA) is a systemic autoimmune disease typically characterized by chronic inflammation and deterioration within the joints. Extra-articular and systemic manifestations can also be present depending on the severity of the disease, and some individuals may experience damage to organs such as the heart, lungs, kidneys, and skin [1].

To date, there are a number of well described treatments available for clinically apparent RA. Of the currently available treatments, conventional disease modifying antirheumatic drugs (DMARDs) and biological DMARDs, also known as biologics or biological immunotherapies, are the most effective for long-term management of RA. However, the effectiveness of these treatments at managing disease progression varies among patients [1], and can be influenced by genetic factors [2,3] and the duration of symptoms prior to the first treatment [4–6]. For example, these factors play a role in why nearly one third of patients do not respond to methotrexate, one of the most common first lines of defense against RA [2,5]. Such interpatient variability in terms of response to medication can interfere with a patient’s ability to achieve remission and/or the desired level of disease activity.

Multiple meta-analyses have used pharmacogenetic studies to summarize how the efficacy, toxicity, and other adverse reactions of conventional DMARDs, such as methotrexate [2,7,8], and biologics [3] such as adalimumab and etanercept can be influenced by genetics. These studies point to polymorphisms in a variety of genes associated with a particular medication’s metabolism, transport, target, and/or mechanism of action as a reason to why response to medication varies among patients. Additionally, the time-frame from start of symptoms to initiation of treatment provides another variable affecting patient response, with a small proposed window of opportunity for a patient to achieve remission. While there is some debate as to the exact time frame of this window [4], there is evidence that treatment within the first 12 weeks of symptoms can lead to lower disease activity scores and remission [9,10]. Other studies indicate a larger window for such outcomes, such as 8 months [11], while the European League Against Rheumatism recommendations for early arthritis treatment indicate a shorter window, with the ideal initiation of treatment beginning within 6 weeks of symptoms [12]. Despite this variability, it is widely accepted that early initiation of treatment is associated with better disease outcomes, and that the time-point at which a patient begins treatment can influence patient response [13].

While the previously mentioned factors interfere with a patient’s ability to achieve remission and/or the desired level of disease activity via physiological mechanisms, they can also interfere with a patient’s ability and/or willingness to adhere to a treatment regimen. Previous examinations of drug retention rates show that patients terminate therapy due to reasons of toxicity and/or lack of efficacy [14–16]. Moreover, initiation of treatment and medication adherence is also influenced by the cost of these therapies. A recent study analyzing the cost of biologic therapies for RA indicated that the average annual cost of treatment with biologics range from a low of around $33,400 for medications like adalimumab and etanercept, to a high of $44,387 for other treatments such as abatacept [17]. While conventional DMARDs (*e.g.* methotrexate) are far less expensive than biologics, interpatient response variability may require that a patient seek methotrexate combination therapy with a biologic, or treatment with a biologic altogether.

Due to the large economic and physiological burden this disease places on its patients, research has become increasingly focused on ways to: 1) better identify those at risk prior to the onset of symptoms and use that information to more accurately predict RA development, and 2) identify and develop effective preventative treatments targeting RA during the pre-clinical phase of the disease and thereby delay the onset of clinical RA [6,18–21].

### Pre-clinical RA

The pre-clinical phase is commonly defined as the stage of the disease in which an individual experiences local or systemic autoimmunity, evidenced by serological abnormalities (*e.g.* high levels of CRP, TNF-□, etc.) and/or autoantibodies (*e.g.* anti-cyclic citrullinated peptide (ACPA), rheumatoid factor (RF), etc.), in the absence of clinical arthritis; many investigators also consider the presence of arthralgia and morning stiffness without clinical arthritis to fall into the pre-clinical phase [13,20,22,23].

Prior to clinical arthritis, initiation and development of autoimmunity is largely influenced by the activation of autoreactive T helper cells (T_h_ cells) from their inactive precursor, naïve CD4^+^ T cells. Once activated, CD4^+^ T cells differentiate into an effector cell phenotype (*e.g.* T_h_1, T_h_2, T_h_17, etc.), each with its own characteristic immune response. T effector cells contribute to inflammation and autoimmunity by coordinating and enhancing the responses of other immune cells. T cell help via T follicular helper cells (T_fh_), for example, plays a critical role in germinal center formation and the production of autoantibodies by B cells [24]. Research has shown that autoantibodies, such as ACPA and RF, can be present in an individual up to ten years before manifestation of arthritic symptoms [25,26], indicating the T_fh_-B cell interactions are occuring long before clinical arthritis. While not all individuals with measurable levels of serum autoantibodies develop clinical arthritis, about 40-60% of individuals who are ACPA positive develop RA within 2-5 years [20,25,27].

Involved in T cell activation are antigen presenting cells (APCs), which present naïve CD4^+^ T cells with two key signals— the presentation of antigen (signal 1) and an interaction between co-stimulatory molecules CD80/86 on the surface of the APCs and CD28 on the surface of CD4^+^ T cells (signal 2). Without both of these signals the naïve CD4^+^ T cells cannot become activated. Due to the naïve CD4^+^ T cells’ dependence on both of these signals, this interaction represents an important target for both prophylactics and treatments of autoimmune diseases [28].

Targeting individuals in the pre-clinical phase of the disease with preventative therapies would provide the earliest initiation of treatment possible, and could halt disease progression prior to significant joint damage. Much of the current research exploring potential prophylactic treatments are mainly focusing on the use of conventional DMARDs, such as hydroxychloroquine (StopRA trial: NCT No. 02603146), and biologics, such as abatacept (APIPPRA study: ISRCTN No. 46017566) and rituximab [29]. Due to the previously mentioned interpatient response variability elicited by this approach, as well as the possible economic and physiological costs of such therapies, it is prudent to explore other prophylactic treatment options--such as alternative, broader spectrum therapies, as opposed to conventional DMARDS or biologics which act through specific, targeted pathways. Furthermore, since the inflammatory load is far less in patients in the pre-clinical phase than in patients experiencing clinical arthritis, it presents an opportunity to potentially use lower-cost alternative therapies that have less adverse reactions.

### Berberine

One potential prophylactic candidate, berberine (BBR), merits further exploration as it has already proved to be of importance for a variety of diseases through successful clinical trials, such as polycystic ovary syndrome [30,31], type II diabetes [32,33], diarrhea-predominant irritable bowel syndrome [34], psoriasis [35], and osteoarthritis [36]. Additionally, current clinical trials are assessing BBR’s ability to help ulcerative colitis (UC) patients maintain remission (NCT No.02962245), to prevent colorectal cancer development in UC patients who are in remission (NCT No. 02365480), and to prevent the recurrence of colorectal adenomas (NCT No. 02226185).

BBR is a plant-derived isoquinoline alkaloid found in the roots, rhizomes and stem bark of plants within a variety of genuses, such as *Berberis* (its namesake), *Mahonia, Hydrastis*, and *Coptis*, among others. The full breadth of botanical sources, as well as the variety of extraction methods, are well-described in a recent review by [37]. Currently, BBR is marketed as a dietary supplement by numerous nutraceutical companies for the treatment of polycystic ovary syndrome, type II diabetes, intestinal inflammation, and as a general anti-inflammatory.

As BBR has been used in human subjects for a number of years, much is already known about its general toxicology and common side effects. Side effects reported after oral administration are considered to be mild (*e.g.* diarrhea, flatulence, abdominal pain, nausea, and constipation), and do not occur in all patients [30,32,38,39]; there were no adverse effects observed on liver and kidney function [32,33,40]. Notably, amelioration of side effects in patients has been reported once dosage was lowered [33]. Additional clinical trials are currently being conducted to gain further insight into absorption mechanisms (NCT no. 03438292), as well as side effects and safety for UC patients in remission who are currently taking BBR as a treatment (NCT No. NCT02365480).

BBR has demonstrated the ability to inhibit the production of a variety of pro-inflammatory cytokines by various immune cells [41,42], having a strong anti-inflammatory effect. It has also been shown to successfully and strongly regulate the inflammatory responses involved in clinically apparent autoimmune diseases *in vivo* such as collagen-induced arthritis [43–46], type I diabetes mellitus [47], UC [48,49], and experimental autoimmune encephalomyelitis [50].

In regard to RA specifically, BBR has been successful at treating clinically apparent collagen induced arthritis (CIA) *in vivo* (the rodent model of RA) through a number of suggested mechanisms: such as **(1)** dendritic cell apoptosis [43], **(2)** interference with MAPK signaling via inhibition of p-ERK, p-38, and p-JNK [44], **(3)** attenuation of T_h_17 activity via inducing cortistatin in the gut [46], **(4)** restoration between balance of T_reg_/T_h_17 cells [45], **(5)** the suppression of Th17 differentiation/proliferation through inhibition of CD169 and RORγt transcription factor, and **(6)** induction of T_reg_ differentiation through aryl hydrocarbon receptor (AhR) activation [51].

Ultimately, many of these mechanisms result in a reduction of both anti-CII autoantibody and inflammatory cytokine production. However, despite evidence that BBR ameliorates *clinically* apparent CIA, and may have prophylactic potential *in vitro* [52] and *in vivo* [53] for other autoimmune pathologies, to date there are no studies involving the use of BBR which explore its prophylactic, *pre-clinical* potential in a CIA mouse model. This is further justified given the substantial population of humans identified as at-risk for RA development and/or with pre-clinical RA (above). Thus, we examine such effects to determine whether or not BBR merits further exploratory analysis as a prophylactic treatment for patients in the pre-clinical phase of RA.

## II. Materials and methods

### General Reagents

DMSO (VWR, Radnor, PA), isoflurane (VetOne, Boise, Idaho), bovine type II collagen in Complete Freund’s Adjuvant (Hooke Labs, Lawrence, MA, USA), 1X PBS, berberine hydrochloride (Sigma Aldrich, St. Louis, MO), ACK lysis buffer (Quality Biological, Gaithersburg, MD), RPMI 1640 supplemented to 2mM L-glutamine, 1% v/v penicillin/streptomycin, 1mM sodium pyruvate, 10mM HEPES (all from ThermoFisher, Waltham, MA) 0.05mM beta-mercaptoethanol (Bio-Rad, Hercules, CA), 10% fetal bovine serum (VWR/Seradigm, Radnor, PA)

### Antibodies

Brilliant Violet 421 anti-mouse CD4 (clone GK1.5), PE anti-mouse CD4 (clone GK1.5), FITC anti-mouse CD3e (clone 145-2C11), FITC anti-mouse CD19 (clone 1D3/CD19), APC anti-mouse CXCR5 (clone L138D7), Alexa Flour 647 anti-mouse FOXP3 (clone MF-14), APC anti-mouse I-A/I-E (clone M5/114.15.2), PE anti-mouse I-A/I-E (clone M5/114.15.2), PE anti-mouse CD80 (clone 16-10A1), APC anti-mouse CD80 (clone 16-10A1), APC anti-mouse CD86 (clone GL-1), PE anti-mouse CD40 (clone 3/23), PE anti-mouse CD25 (clone 3C7), FITC anti-mouse CD25 (clone 3C7), APC anti-mouse CD154 (clone MR1), APC anti-mouse CD28 (clone 37.51), FITC anti-mouse CD28 (clone E18) and recommended isotype controls (all from BioLegend, San Diego, CA).

### Mice

DBA/1J mice (6 weeks old) were purchased from Jackson Laboratories (Bar Harbor, ME). Animals were acclimated to the housing facilities for one week prior to starting experiments. All procedures were approved by the Institutional Animal Care and Use Committee (protocol # 1801BD-NP-M-21) and performed at the University of Northern Colorado in accordance with institutional and international guidelines. Mice were divided into four groups: Control (no CIA induction, no treatment), CIA (positive control), PBS+DMSO (volume-matched vehicle control), and BBR (berberine treatment, 1mg/kg per day). Before commencing the full experiment involving cellular analyses a pilot study was performed to determine the efficacy of the CIA model (n=3 per group). Treatments were administered via I.P. injections 5 times per week, with 5 days on and 2 days off. With the full study (n=15 per group) five mice from each group were euthanized on day 14 (pre-clinical stage), with 10 mice from each group being euthanized on day 28.

### CIA induction and assessment

An emulsion of bovine type II collagen and Complete Freund’s Adjuvant (Hooke Labs, Lawrence, MA, USA) was injected according to manufacturer’s instructions. Briefly, mice were anesthetized with isoflurane and 0.05 mL injections of the emulsion were delivered subcutaneously approximately 1 inch distal from the base of the tail, in the space between the ventral and lateral tail veins. The needle was left inserted for 5 to 10 seconds post-injection to help prevent any possible leakage of the emulsion. This procedure was repeated with all mice in the CIA, CIA+BBR, and CIA+PBS groups, and was considered day 0 of the experiment. For mice undergoing full observation through day 28, on day 18 a booster injection of bovine type II collagen and Incomplete Freund’s Adjuvant emulsion was given. The emulsion was injected subcutaneously on the same side of tail as, although slightly cranial to, the site of the initial immunization injection. On day 28, mice were evaluated for the presence of arthritis, and scored on a scale of 0-16 as follows (per manufacturers’ recommendation) : 0 = normal paw, 1 = one toe inflamed and swollen, 2 = more than one toe, but not entire paw inflamed and swollen OR mild swelling of entire paw without ankle swelling, 3 = entire paw inflamed and swollen (inclusion of ankle swelling), 4 = severely inflamed and swollen OR ankylosed paw; all paws were assessed for a total possible score of 16.

### ELISA

Blood samples were collected immediately following euthanasia of mice on day 14 and 28, and serum concentrations of anti-collagen type II (anti-CII) total IgG (catalog # 1012T), anti-CII IgG1 (catalog # 20321T), and anti-CII IgG2a (catalog # 20322T) were measured by ELISA (Chondrex, Redmond, WA, USA) according to the manufacturer’s instructions. Briefly, plates pre-coated with bovine collagen II, were washed with 1X PBS containing 0.05% polysorbate-20 and blocked (PBS/0.05 % bovine serum albumin, latter from VWR) for 2h at room temperature. Blocking buffer was added at 100 μl/per well and incubated for 1 hr at room temperature. Standard dilutions and serum samples (diluted 1:1000) were added at 100 μl/per well and incubated at room temperature for 2 hr. Plates were washed and secondary antibody solution for total IgG or specific IgG subtypes was added at 100 μl/per well and incubated at room temperature for 1 hr. Plates were washed and TMB solution was added at 100 μl/per well for 15 minutes at room temperature. Stop solution (2N sulfuric acid) was added at 50 μl/per well and optical densities were taken at 450 nm using a microplate reader.

### Flow cytometry

Single cell suspensions were made from spleens, inguinal lymph nodes (LNs), and axillary LNs of euthanized mice and stained with fluorescent antibodies specific for cell lineage markers: CD4^+^CD3^+^Foxp3^-^T helper (T_h_) cells, CXCR5^+^ T follicular helper (Tfh) cells, CD4^+^Foxp3^+^ T regulatory cells (T_regs_), and CD19^+^ B cells were measured at days 14 and day 28. Spleen and LN cells were also stained with fluorescent antibodies specific for co-stimulatory molecules involved in T cell and B cell activation and differentiation: CD40L and CD28 on CD4^+^CD3^+^Foxp3^-^ T_h_ cells, CD25 on CD4^+^Foxp3^+^ T_regs_, and MHC class II, CD40 and CD80/86 on CD19^+^ B cells were measured at day 14 and day 28.

### Statistical analysis

Shapiro-Wilk was used to test for normality, and the distribution was not normal. Chi-square (χ^2^) was used to compare absolute arthritic incidence (score ≤ 2) among control and treatment groups. For all other comparisons, due to violation of normality, the Kruskal-Wallis test with Dunn’s multiple comparisons and/or Mann-Whitney U test were used. All tests had an α = 0.05.

## III. Results

### Berberine treatment delays onset of CIA

To assess BBR’s ability to delay the onset of clinical CIA, mice were observed daily for signs of redness and joint swelling as an indication of arthritis development, and severity of arthritis was scored on a scale of 0-16 as previously described. When mice were euthanized on day 28, we observed a significant reduction in absolute incidence of arthritis in the BBR group compared to the CIA and PBS controls (Fig. 1A). About 90% of mice in the CIA control group developed arthritis, compared to 80% in the PBS control group and 40% in the BBR group. In mice who developed arthritis, however, there was a trend but no significant difference in severity (Fig. 1B).

**Figure 1.**
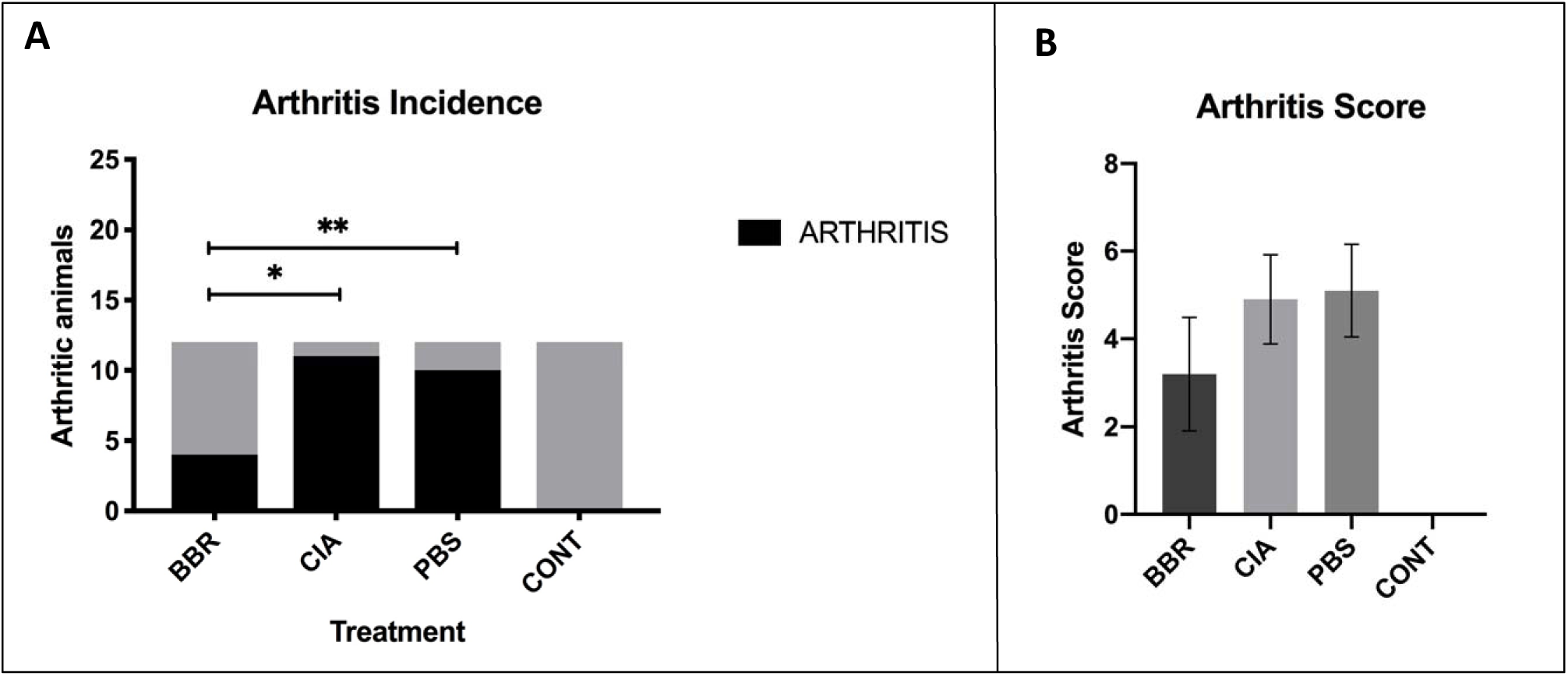
Assessment of CIA in DBA/1J Mice in the Context of BBR Treatment. A. Absolute incidence of arthritis (proportions of animals with score ≤ 2) among treatment groups at day 28 compared using χ^2^ (n=12 per group, *p<0.05). Incidence proportions were BBR = 40%, CIA = 90%, PBS = 80%, and CONT = 0%. B. Arthritis score, on a scale of 0-16 per manufacturer protocol (as described in Methods), of mice at day 28 treated with BBR (1 mg/kg/day), volume-matched 1X PBS with .01% DMSO (PBS vehicle control), or no treatment (CIA Control). Multiple comparisons conducted using the Kruskal-Wallis test with Dunn’s multiple comparisons (n=12 per group).

### The effect of berberine on circulating anti-CII IgG in the CIA model

To determine if BBR prophylactic treatment reduces autoantibody production, serum concentrations of anti-CII total IgG, anti-CII IgG1, and anti-CII IgG2a autoantibodies were measured at the day 28 endpoint. BBR group saw significantly reduced serum concentrations of anti-CII IgG2a and anti-CII total compared to both CIA and PBS controls, although there was no significant difference in anti-CII IgG1 in BBR mice compared to CIA control mice (Fig. 2A). To further examine if the aforementioned results were an artifact of including both arthritic and non-arthritic mice in the dataset, comparisons of just arthritic mice were performed. In this comparison, levels of anti-CII IgG2a among arthritic mice in the BBR group remained significantly reduced compared to CIA and PBS controls (Fig. 2B). When comparing anti-CII IgG levels between arthritic and non-arthritic mice within the BBR group specifically, however, anti-CI IgG1, IgG2a and total IgG were all significantly reduced in the non-arthritic mice compared to those who developed arthritis (Fig. 2C). Additionally, there appeared to be a vehicle-specific effect on circulating anti-CII IgG in which the administration of PBS with 0.01% DMSO elicited elevated levels of anti-CII IgG1 and total IgG in vehicle control mice (Fig. 2A-B).

**Figure 2.**
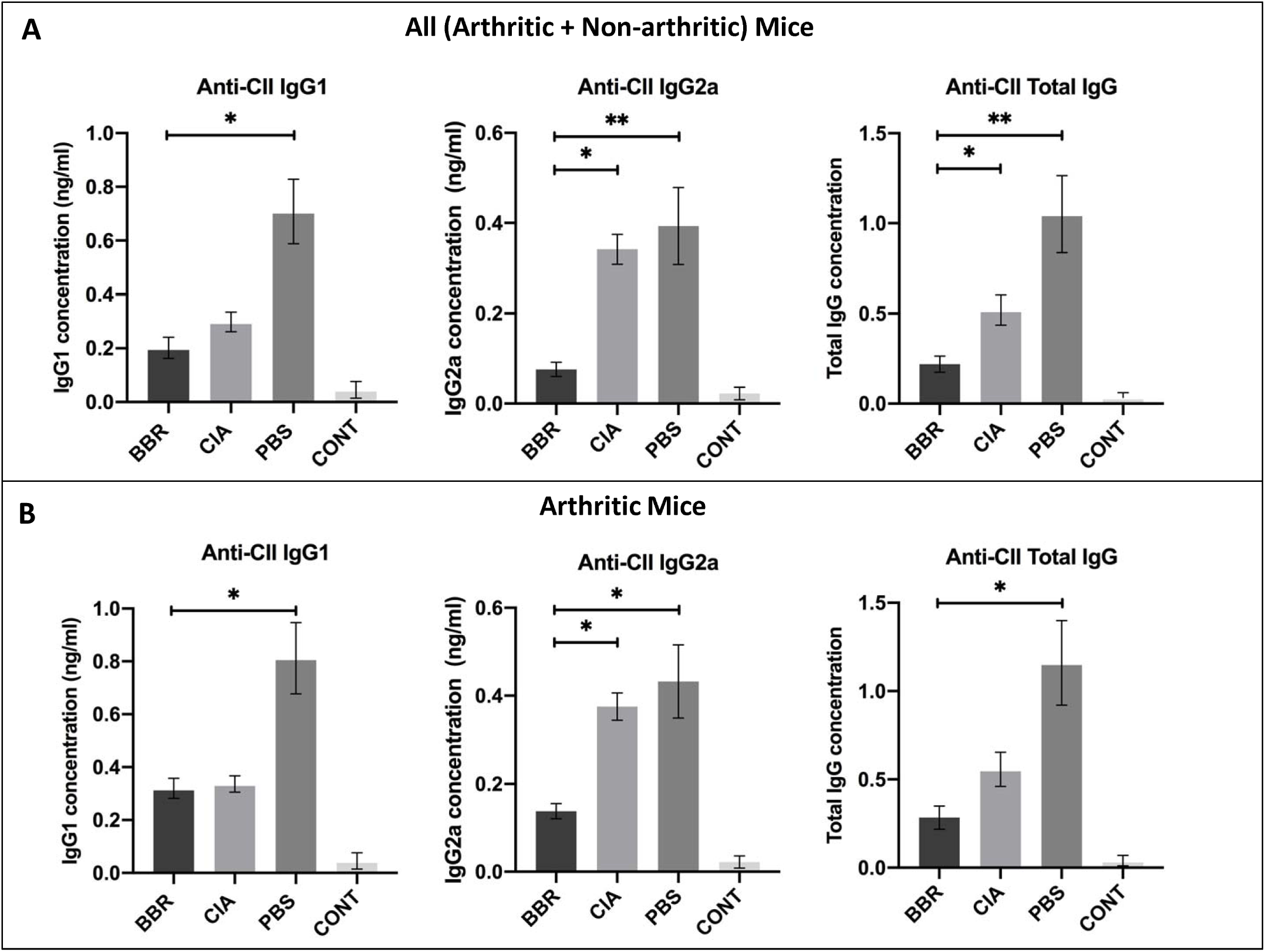
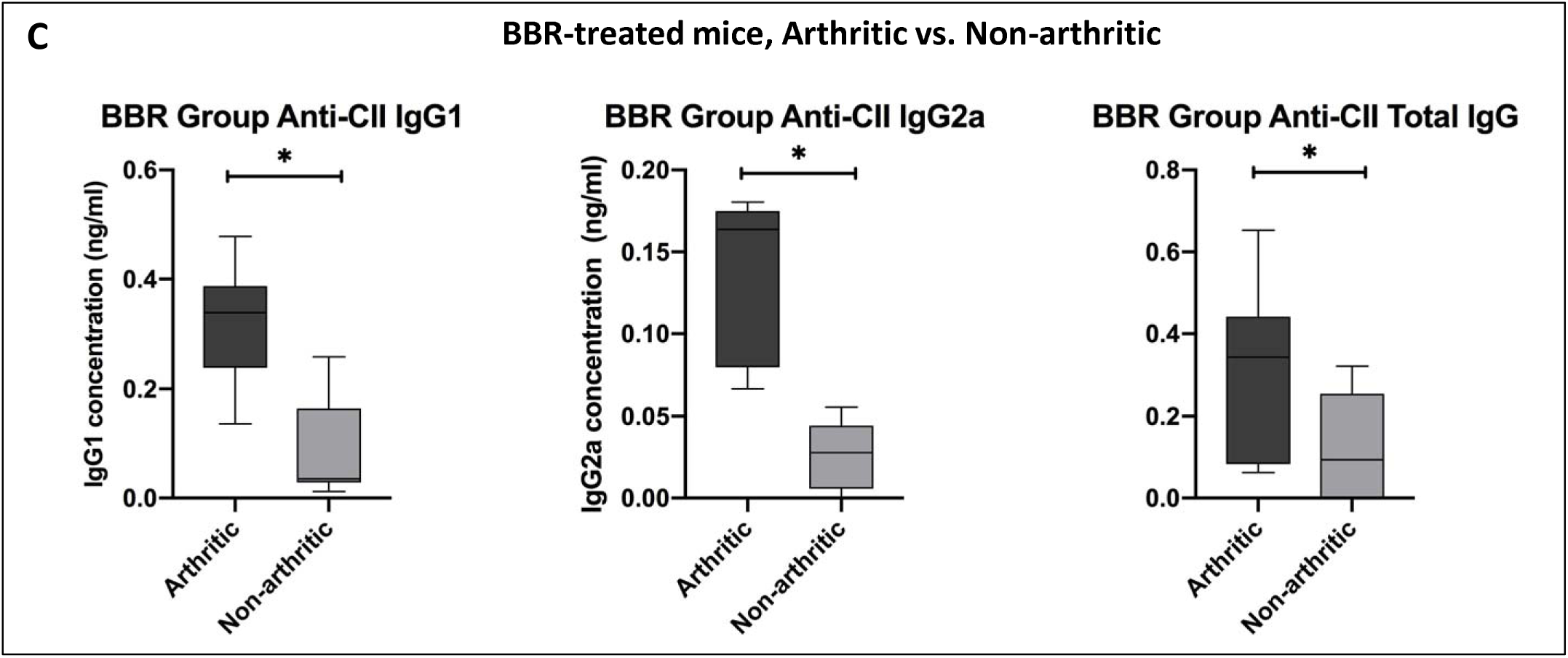
The effect of berberine on circulating anti-CII IgG in the CIA model. **A.** Anti-CII IgG1, IgG2a, and total IgG at day 28 among all mice (arthritic and non-arthritic) within BBR, PBS (vehicle control), CIA (no treatment control), and non-CIA control animals (n=12 per group). **B.** Anti-CII IgG levels at day 28 compared among arthritic mice only (BBR n=5; PBS n=10; CIA n=11). Statistical comparisons made with the Kruskal-Wallis test with Dunn’s multiple comparisons (*p<0.05). **C.** Anti-CII IgG levels at day 28 compared among arthritic vs. non-arthritic mice who were treated with BBR. Statistical comparisons made with the Mann-Whitney U test (*p<0.05).

### Key CD4^+^T cell population characteristics in response to berberine treatment

On day 14, we observed a significant reduction in populations of both CD4^+^T cells and CXCR5^+^T_fh_cells in the LNs and spleen of BBR-treated mice (Fig. 3A-B), as well as a reduction in the percentage of CD28^+^ and CD154^+^ CD4^+^T cells in the spleen and LNs of BBR-treated mice (Fig. 3A-B). By the day 28 experimental endpoint we continued to observe a significant reduction in CD4^+^T cells and CXCR5^+^T_fh_ cells in the spleen and LNs of BBR-treated mice (Fig. 3C-D), however, there was no significant difference in the percentage of CD28^+^ and CD154^+^ CD4^+^T cells between treatment groups (Fig 3C-D).

**Figure 3.**
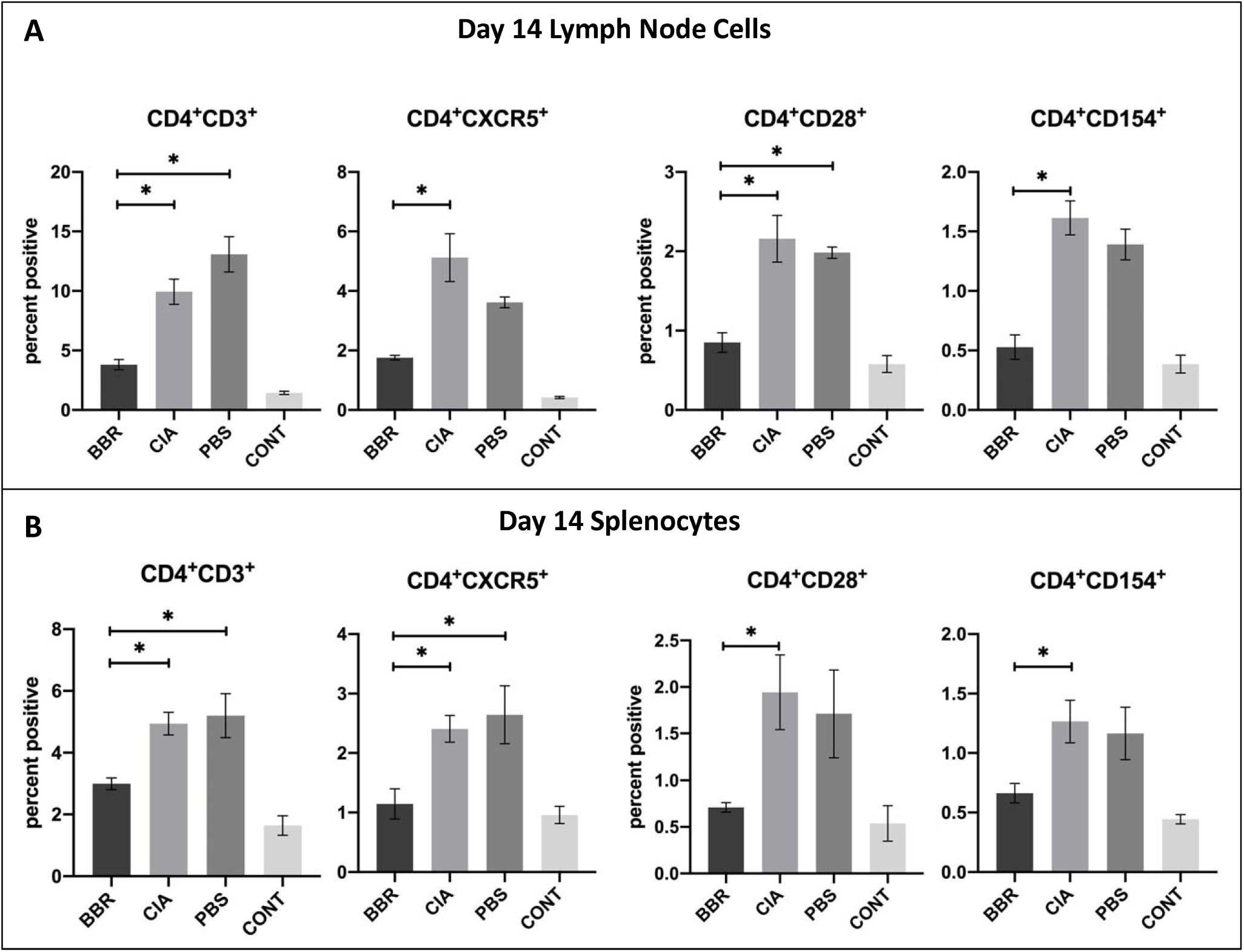
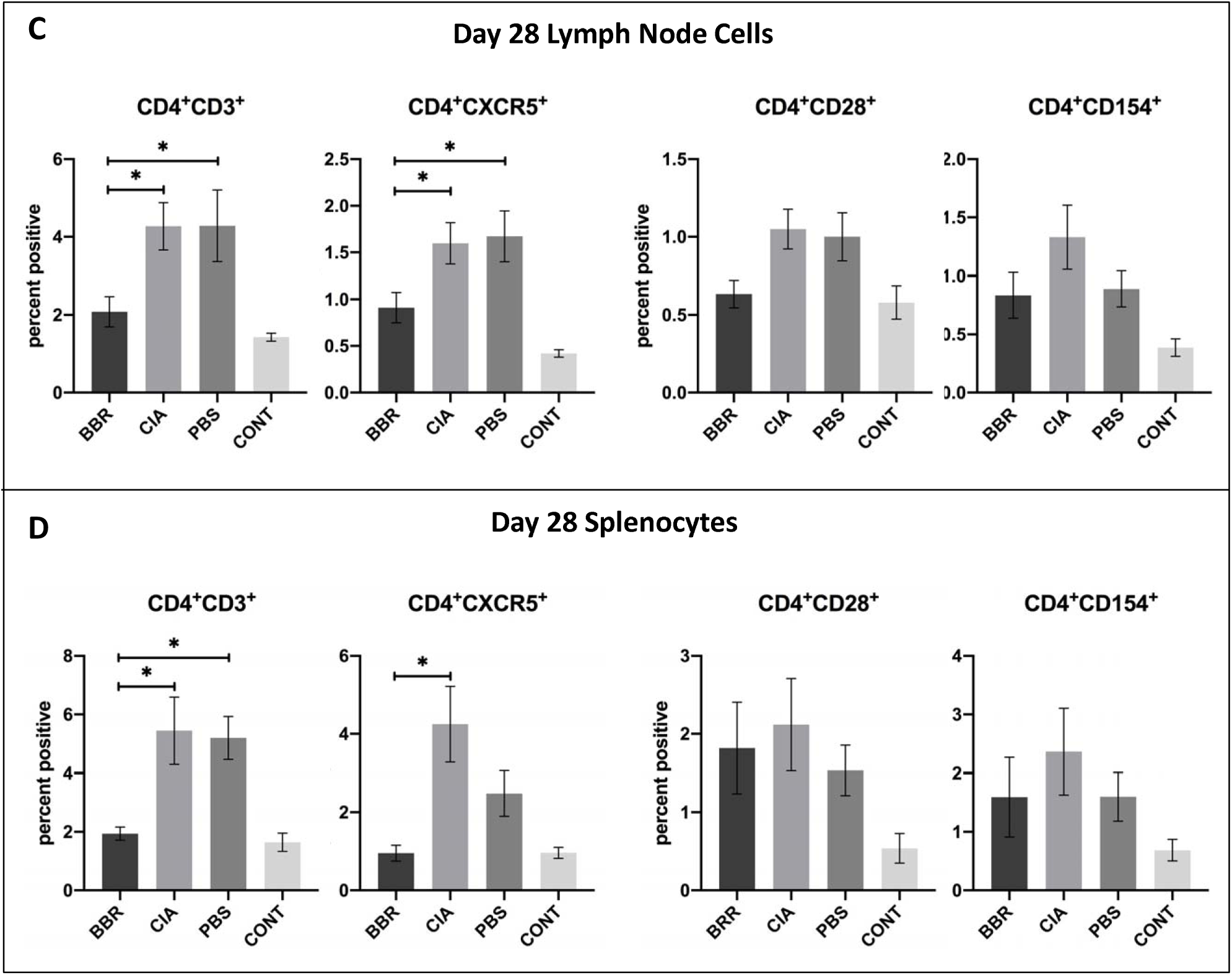
CD4^+^ T cell populations during pre-clinical CIA (day 14) and at final day 28 endpoint. Cells compared were from the CD4^+^ T cell population of LN and spleen with further investigation into CD4^+^ T cell populations expressing specific cell-surface markers. Shown are populations of CD4^+^ T, CXCR5^+^ T_fh_, CD154^+^CD4^+^ T cells, and CD28^+^CD4^+^ T cells of the LN (**A.**) and spleen (**B.**) at the day 14 endpoint (n=5 per group), and of the LN (**C.**) and spleen (**D.**) at the day 28 experimental endpoint (n=10 per group). Statistical comparisons made with the Kruskal-Wallis test with Dunn’s multiple comparisons (*p<0.05).

### Berberine treatment leads to increased proportion of FOXP3^+^CD25^+^CD4^+^ T cells

To examine BBRs effect on T_reg_ populations, cells from the CD4^+^ CD25^+^ T population of LN or spleen were measured for the presence of the definitive T_reg_ transcription factor FOXP3. Out of this subset of cells, we observed an increased ratio of FOXP3^+^: FOXP3^-^ cells in the spleen and LNs of BBR-treated mice during the pre-clinical phase of CIA (day 14 endpoint) (Fig 4A). At the day 28 endpoint, BBR-treated mice had a significantly increased ratio of FOXP3^+^: FOXP3^-^ cells in the LNs, but not the spleen (Fig. 4B). In order to determine whether or not the previously mentioned results were an artifact of including both arthritic and non-arthritic mice in the analysis, we compared this ratio between mice in the BBR group who developed arthritis and the mice who did not. There was no significant difference in the day 28 splenic FOXP3^+^: FOXP3^-^ T cell ratio between arthritic and non-arthritic mice in the BBR group. However, all non-arthritic BBR treated mice had a larger percentage of FOXP3^+^ cells compared to FOXP3^-^ cells (ratio of >1) except for one outlier, whereas all arthritic BBR-treated mice had a smaller percentage of FOXP3^+^ cells compared to FOXP3^-^ cells (ratio of <1) (Fig. 4C).

**Figure 4.**
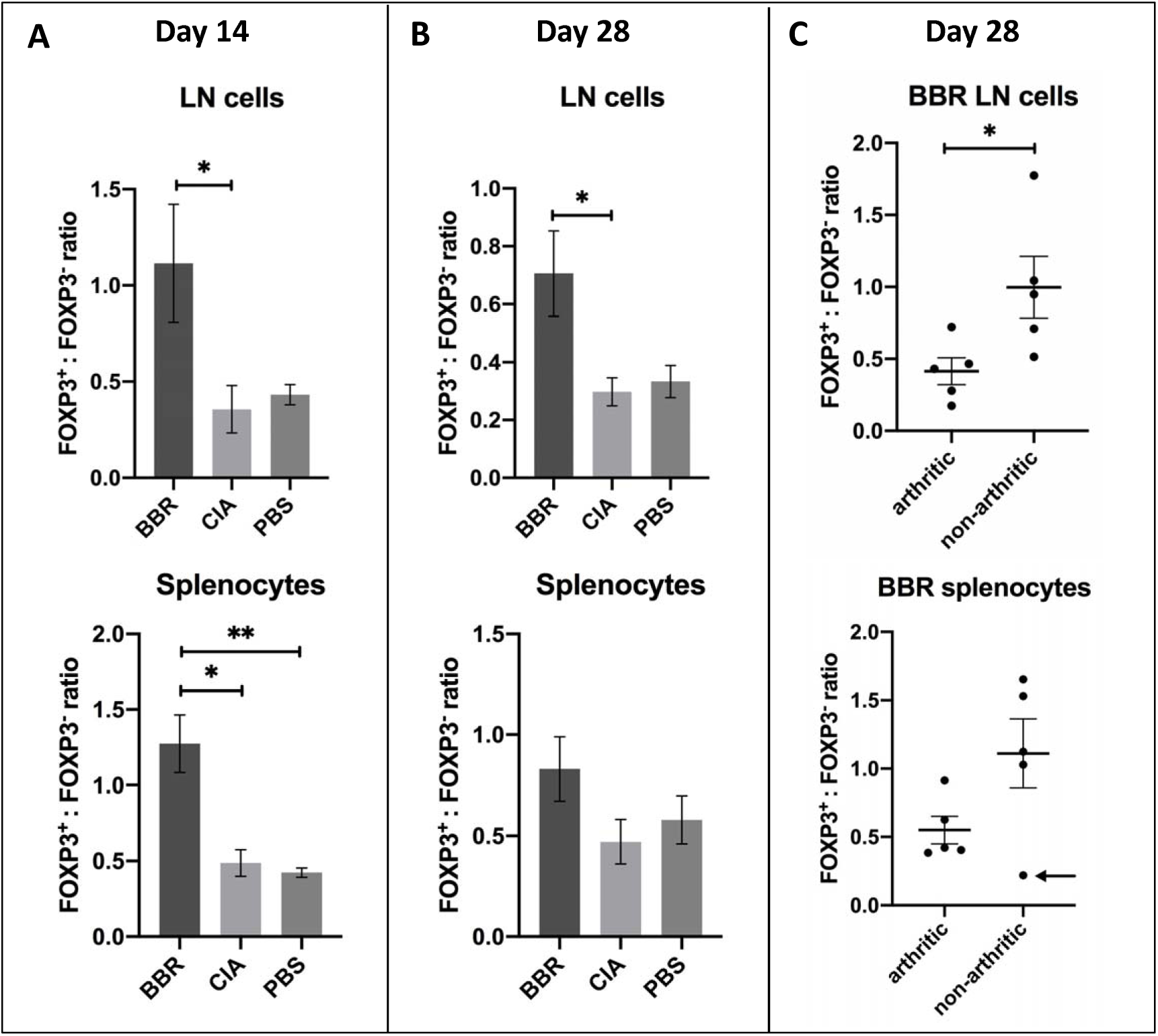
Berberine induces T_reg_ expansion in lymphoid tissue during CIA induction. Cells compared were from the CD4^+^ CD25^+^ T_h_ population of LN or spleen with further interrogation of the definitive T_reg_ transcription factor FOXP3. **A.** The FOXP3^+^:FOXP3^-^ ratio during the pre-clinical phase of arthritis (day 14) (n=5 per group, *p<0.05). **B.** The FOXP3^+^:FOXP3^-^ ratio at the day 28 experimental endpoint (n=10 per group, *p<0.05). Ratios compared using the Kruskal-Wallis test with Dunn’s multiple comparisons.

### Key CD19^+^B cell population characteristics in response to berberine treatment

In regards to berberine’s effect on B cells of the spleen and LNs during CIA development, there was no significant difference in CD19^+^ B cell populations or CD19^+^ cell populations expressing specific cell-surface markers MHC II, CD40, and CD80/86 between BBR-treated and the CIA control mice (Fig. 5A-B). This was despite trends in total B cell reduction, as well as reduction in CD80^+^ and CD86^+^ B cells in both LNs and spleen, and reduction of MHC II^+^ B cells in LNs of BBR-treated mice. However, we did observe a vehicle-specific effect similar to that seen in the anti-CII IgG data. There was a significantly higher CD19^+^ B cell population and percentage of MHC II^+^CD19^+^ B cells in both the spleen and LNs of the PBS control mice compared to the BBR-treated mice. Additionally, this effect was also seen with splenic CD19^+^CD80/86^+^ B cells.

**Figure 5.**
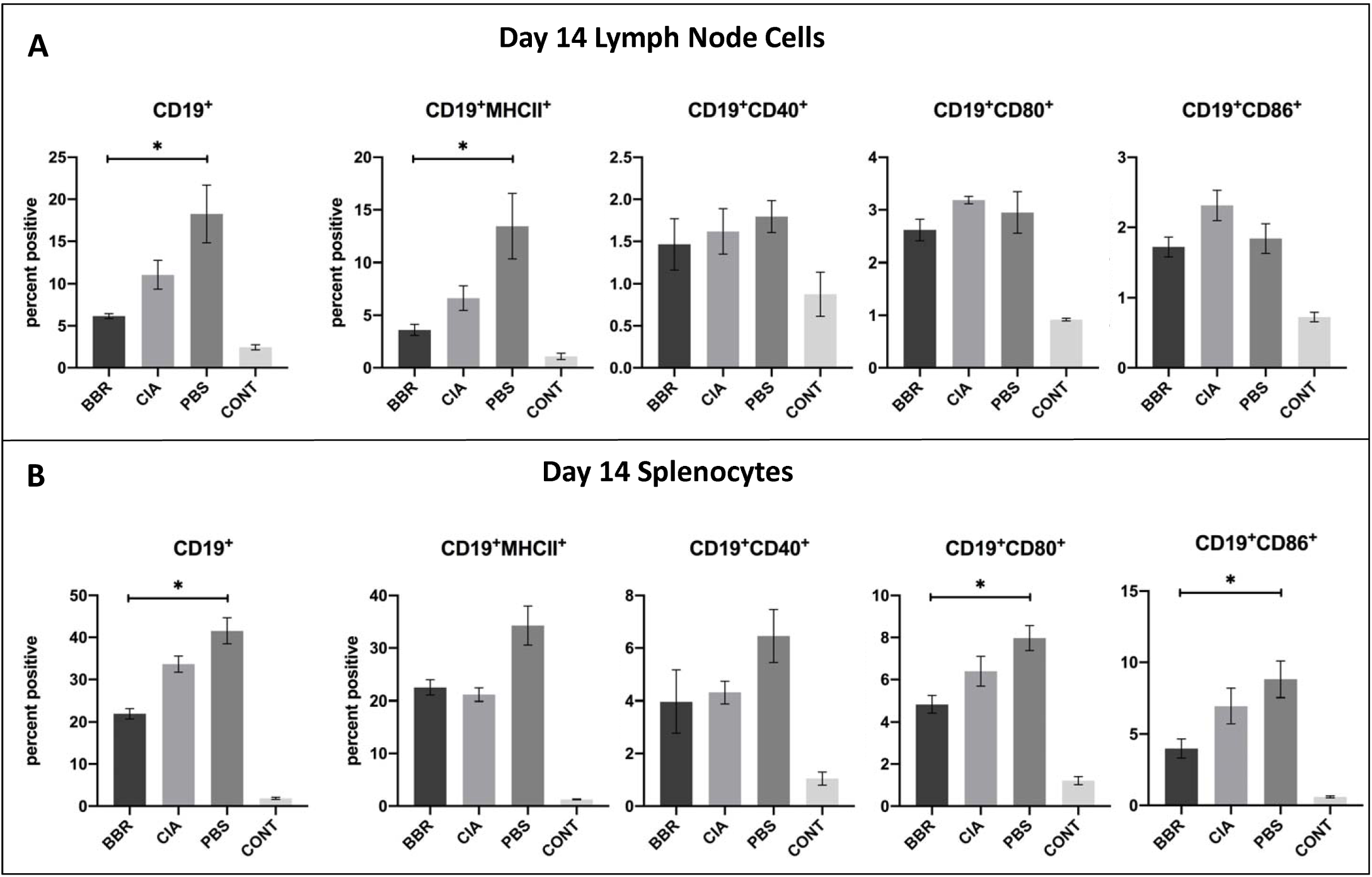
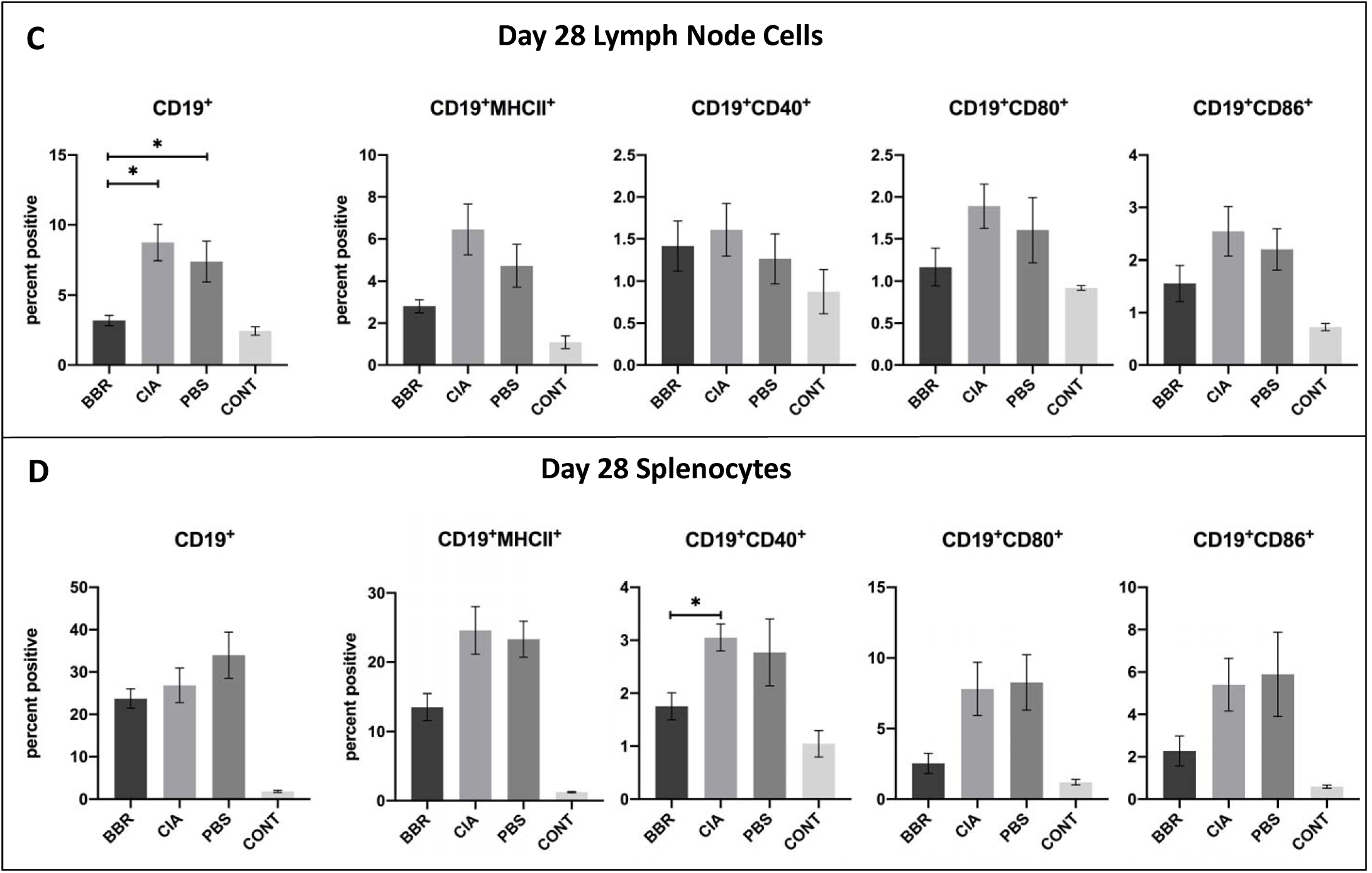
CD19^+^ B cell populations during pre-clinical CIA (day 14) and at final day 28 endpoint. Cells compared were from the CD19^+^ B cell population of LN and spleen with further investigation into CD19^+^ cell populations expressing specific cell-surface markers. Shown are populations of CD19^+^ B cells, MHCII^+^CD19^+^ B cells, CD40^+^CD19^+^ B cells, CD80^+^CD19^+^ B cells, and CD86^+^CD19^+^ B cells of the LN (**A.**) and spleen (**B.**) at the day 14 endpoint (n=5), and of the LN (**C.**) and spleen (**D.**) at the day 28 experimental endpoint (n=10). Statistical comparisons made with the Kruskal-Wallis test with Dunn’s multiple comparisons (*p<0.05).

By day 28 we observed a significant reduction in CD19^+^B cells in the LNs, but not spleen, of BBR treated mice (Fig. 5C-D). While there was a trend there was no significant reduction in CD19^+^ cell populations expressing specific cell-surface markers MHC II, CD40, and CD80/86 in the spleen and LNs of BBR-treated mice, nor CD40 in the LNs of BBR-treated mice. Unexpectedly, and inconsistent with the rest of the data, splenic populations of CD40^+^CD19^+^ B cells appeared to be significantly reduced in BBR-treated mice.

## IV. Discussion

Our results show that BBR treatment during the pre-clinical phase of CIA delayed the onset of CIA in DBA/1J mice, although mice in the BBR group who developed arthritis did not experience a significant decrease in clinical arthritis score compared to the CIA and PBS controls. Based on our results and evidence from other studies, it is likely that this protective effect was directly mediated through effector CD4^+^ T helper cell suppression, which subsequently influenced activation, proliferation, and autoantibody production by B cells.

Our hypothesis that BBR is exerting its effect via CD4^+^ T helper cell suppression is supported through the observation that BBR-treated mice had reduced populations of CD4^+^T cells; while this population included all T cell subsets expressing CD4 (both T_h_ and T_reg_), we observed a higher percentage of FOXP3^+^CD25^+^CD4^+^ T cells (representative of T_reg_) compared to a decrease in CD4^+^CXCR5^+^T_fh_ cell populations, as well as the reduced percentage of CD28^+^CD4^+^ and CD154^+^CD4^+^ T cells during CIA development (day 14). Although this reduction in CD28^+^CD4^+^ and CD154^+^CD4^+^ T cells was not observed at the day 28 point, BBR-treated mice still maintained lower overall populations of CD4^+^T cells and CXCR5^+^T_fh_ cells, as well as a higher percentage of T_reg_ within the total CD4^+^ T cell population.

A previous study by Moschovakis *et al.* (2017) [54] examining the role of CXCR5^+^ T_fh_ cells in RA showed that T cell-specific CXCR5 deficiency prevented RA development. Furthermore, an *in vivo* CIA study involving the use of hydroxychloroquine (HCQ) as a prophylactic (administered from day 0 of the experiment) noted a reduction in T_fh_ cells, which corresponded to a decrease in both incidence and arthritis score in HCQ-treated mice [55]. This same study also observed higher concentrations of circulating T_fh_ cells in the blood of RA patients compared to healthy individuals. Similar to this evidence, it is possible that the reduced populations of CXCR5^+^T_fh_ cells seen in the BBR group compared to the CIA and PBS controls contributed to lower incidence of arthritis as well as the reduced generation of anti-CII total IgG and subtypes, as CXCR5^+^T_fh_ cells play a critical role in germinal center formation, B cell affinity maturation and autoantibody production [24]. As such, we propose the mediation of CXCR5^+^T_fh_ cell proliferation as a novel function of BBR, and we are unaware of any studies to date that specifically look at the effect of BBR on CXCR5^+^T_fh_ cell populations.

Additionally, the reduced percentage of CD28^+^CD4^+^ and CD154^+^CD4^+^ T cells during CIA development (day 14) in BBR-treated mice could be indicative of reduced activation and proliferation of CD4^+^T cells, thereby resulting in the lower CD4^+^ T cell populations observed in the BBR-treated group. Furthermore, in preliminary *in vitro* experiments conducted prior to commencing the full length CIA model (data not published), BBR treatment (10μM) of CD4^+^ T cells during activation with anti-CD3 (plate-bound, 5μg/mL overnight before culture) and soluble anti-CD28 (soluble, 2μg/mL) for 5 days significantly reduced populations of CD4^+^ T cells and expression of CD28 on CD4^+^ T cells (Supplementary Figure 1A), and a trend of reduction in CD154 expression on CD4^+^ T cells. We observed this in addition to a decreased concentration of IL-2 in cell supernatants (Supplementary Figure 1B), both canonical indicators of Th1 activation and autocrine signaling. The blockade to CD4^+^ T cell co-stimulation has proven to be an effective RA treatment and is the mechanism of action of abatacept, a biological immunotherapy used to treat clinically apparent RA [1,21]. CD28-CD80/86 interaction is an important therapeutic target as CD28 ligation leads not only to increased T cell proliferation and activation, but also to increased CD154 expression [28]; CD154 is a crucial ligand involved in the activation of B cells and other APCs.

BBR’s protective effect against CIA development is also likely mediated through it’s alteration of the FOXP3^+^: FOXP3^-^ CD4^+^ T cell ratio. With the exception of the day 28 splenocytes whose data was skewed by one outlier, the BBR group saw a significantly higher proportion of FOXP3^+^CD25^+^CD4^+^ T cells (T_reg_) compared to CIA and PBS controls. Thus, while BBR treatment resulted in lower overall CD4^+^ T cell populations, a higher percentage of cells within that reduced population were T_reg_. Previous studies corroborate the protective effect of T_reg_ on CIA development; adoptive transfer of CD25^+^ T_reg_ slowed disease progression in a CIA model [56], and the depletion of CD25^+^ T_reg_ prior to immunization with bovine type II collagen (used to induce CIA) exacerbated arthritis [57].

In reference to BBR and CIA specifically, prior studies using BBR to ameliorate clinically apparent CIA resulted in a suppression of T_h_17 activity alongside the activation/proliferation of T_reg_, thereby resulting in an increased T_reg_/T_h_17 ratio in BBR treated mice [45,51]. While a study by Yue *et al.* (2017) [46] provides opposing evidence in which BBR did not appear to have significant effect on the frequency of T_reg_ in a CIA model despite seeing amelioration of clinically apparent CIA, their particular model used PBMCs to assess T_reg_ population and FOXP3 expression, as opposed to our study which used splenocytes and draining LN cells. Moreover, animal models of autoimmune diseases other than RA/CIA indicate a protective role for BBR through the increase in T_reg_ population relative to pro-inflammatory cell types. Previous research examining the efficacy of BBR as a DSS-induced UC treatment, for example, reported that BBR improved the T_reg_/T_h_17 balance [58]. One possible mechanism for our observed increase in the proportion of T_reg_ is via AhR signaling; two studies involving the use of other alkaloids, sinomenine [59] and norisoboldine [60], ameliorated CIA and increased T_reg_ populations in an AhR-dependent manner; BBR has also been shown to activate AhR [61–63]. Additionally, one study examining the protective effect of keratinocyte growth factor on CIA development reported that there were increased percentages of T_reg_ in animals who experienced delayed onset [64], providing evidence for the protective role of T_reg_ in delaying the onset of CIA.

In regard to B cell-specific responses to BBR, during CIA development (day 14 endpoint) the BBR-treated mice in our study did not see a significant reduction in overall CD19^+^ B cell populations or populations of co-stimulatory molecule expressing CD19^+^ B cells compared to the CIA control. However, by day 28 we observed significant reduction in CD19^+^ B cell populations in the draining lymph nodes of BBR-treated mice, as well as a reduction in anti-CII IgG2a and total IgG. As such, we propose that the reduction in day 28 lymphatic B cell populations and subsequent lowering of anti-CII autoantibody production is largely due to BBR interfering with the T-cell mediated activation of B cells via T cell suppression, thereby contributing to decreased B cell activation. This interference could be due not only to the decreased CD4^+^CXCR5^+^ T_fh_ cell populations and enhanced proportion of T_reg_ seen throughout the experiment in BBR-treated mice, but also the decrease in CD28^+^CD4^+^ and CD154^+^CD4^+^ T cells seen during disease development (day 14 endpoint). Both CD28-CD80/86 and CD154-CD40 interactions play an important role in B cell activation and proliferation, and CD154-CD40 ligation specifically provides key signaling for thymus-dependant humoral immunity responses, such as the isotype class-switching and affinity maturation required to generate high affinity anti-CII IgG autoantibodies [65–67]. Furthermore, previous research has demonstrated that the disruption of CD28-CD80/86 and CD154-CD40 interactions results in reduced anti-CII autoantibody titres, prevention of disease development, and/or amelioration of disease in CIA and other autoimmune arthritis models [68–72]. In other words, pro-inflammatory T cell development and activation is inhibited by BBR early, which leads to a later reduction in B cells reactive to CII stimulus, and this timing fits with classical T cell-mediated B cell activity.

While there was no significant difference in anti-CII IgG1 observed between the BBR group and CIA control, we did observe a significant reduction in anti-CII IgG2a. Moreover, BBR-treated mice who experienced a delay in onset (non-arthritic by day 28) had significantly lower concentrations of anti-CII IgG1, anti-CII IgG2a and anti-CII total IgG compared to arthritic mice. In CIA the IgG subtype that is thought to play the biggest direct role in inflammation and joint destruction is anti-CII IgG2a, which predominantly activates the complement cascade, although it can also bind Fcγ receptors (Fcγ R) on FcγR-bearing immune cells. High concentrations of anti-CII IgG1 are also typically present, however IgG1 more readily binds to and activates FcγR-bearing immune cells and has a lower affinity for activating complement compared to IgG2a [73,74]. The important role of complement activation in CIA pathology is supported by studies that demonstrated amelioration of CIA in response to complement deficiency [75] and that C5-deficient mice were resistant to CIA development [76]. As IgG2a is a strong activator of complement, IgG2a serum concentration has been shown to correlate to the degree of inflammation as well as cartilage and bone destruction in CIA models [77], and reduced serum concentrations of IgG2a were associated with delayed onset and reduced frequency of arthritis incidence [78,79]. However a notable difference with our study is that while we observed significantly lower concentrations of IgG2a in BBR-treated mice compared to CIA and PBS controls, we did not see any significant difference in degree of observable inflammation (arthritis scores). Additionally, as previous studies involving the use of BBR to treat clinically apparent CIA reported a significant reduction in anti-CII IgG1 in BBR-treated mice compared to both CIA and PBS controls [43,44], the lack of significant anti-CII IgG1 reduction in the BBR group compared to the CIA control in our own study was unexpected. It is notable, however, that when comparing arthritic and non-arthritic mice within the BBR group alone, the non-arthritic mice had significantly lower concentrations of both anti-CII IgG2a and anti-CII IgG1, indicating that the observed reduced incidence of arthritis is likely in part due to a reduction in circulating autoantibodies, as seen in other studies [78,79].

One major unexpected result regarding anti-CII autoantibody production involved a vehicle-specific effect in which the PBS control group saw the highest increase in autoantibody production in compared to the CIA control and BBR group. Our solution of BBR dissolved in PBS and 0.01% DMSO was modeled in part after a previous CIA study that used BBR dissolved in a PBS/DMSO solution containing a slightly greater concentration DMSO than our own [44]; this previous study did not report elevated levels of anti-CII total IgG or anti-CII IgG subtypes in PBS control groups. However, DMSO has demonstrated the ability to stimulate antibody production from hybridoma cells, which are myeloma-B cell hybrids commonly used to generate large quantities of monoclonal antibodies in research and industry settings [80]. In light of this, it is possible we witnessed a B cell-specific response to the presence of DMSO.

In addition to this unexpected vehicle-specific effect, our model also faced limitations. One major limitation was the final day 28 endpoint; Prolonging the final endpoint past day 28 would provide more insight into preventative capabilities of BBR. Due to the fact that our non-arthritic mice continued to have suppressed populations of CD4^+^ T helper cells and CXCR5^+^ T_fh_ cells, higher relative percentages of T_reg_, and lower concentrations of circulating autoantibodies by day 28, we hypothesize that it is likely BBR treatment would at least continue to delay CIA development to a certain point. However, it is not known whether BBR would entirely prevent CIA development in those mice who remained non-arthritic by day 28, or if they would eventually develop symptoms clinical arthritis at a later date. This model is further limited in that it assumes a mouse would be able to absorb the I.P. administered dose via oral administration, which is the preferred route of administration for human patients taking BBR dietary supplements. While estimates vary, it is widely known that BBR has an extremely low oral bioavailability (<1%) [81–83]. Thus, this model is not entirely reflective of how a human patient would ideally receive and BBR as a treatment, nor of how a patient would absorb and distribute BBR as an orally delivered treatment.

In conclusion, BBR is likely having protective effects against CIA development by directly suppressing CD4^+^ T helper cell activity, thus having an indirect effect on B cell activation and autoantibody production. These T cell suppressive effects are evidenced by a lower percentage of co-stimulatory molecule-expressing CD4^+^ T cells during CIA development (day 14), as well as lower populations of CD4^+^ T cells (including CXCR5^+^ T_fh_ cells), and higher percentages of T_reg_ in BBR-treated mice throughout the experiment. These suppressive effects are not reflected in the percentage of B cells expressing key co-stimulatory molecules, although populations of CD19^+^ B cells were lower in the draining lymph nodes of BBR-treated mice by day 28, indicating that reduced B cell proliferation and anti-CII auto-antibody production is likely due to decreased interaction with activated T_h_ cells. In the future, it is important to repeat this experiment with a later endpoint to better determine the duration of BBR’s protective effects, as well as more closely examine BBR’s influence on CXCR5^+^ T_fh_ cells and germinal center formation. As the formation of germinal centers, plasma cells, and memory B cells are crucial to our adaptive immunological memory, BBR’s suppressive effect on CXCR5^+^ T_fh_ cell populations raises concern that prolonged use could potentially impact a patient’s ability to mount effective secondary immune responses.

## FUNDING

This work has been supported by the University of Colorado NHS Student Research Fund and Graduate Student Association Grants

## ACKNOWLEDGMENTS

N/A

## DISCLOSURE

The authors declare no conflict of interest.

---

**Supplementary Figure 1.**
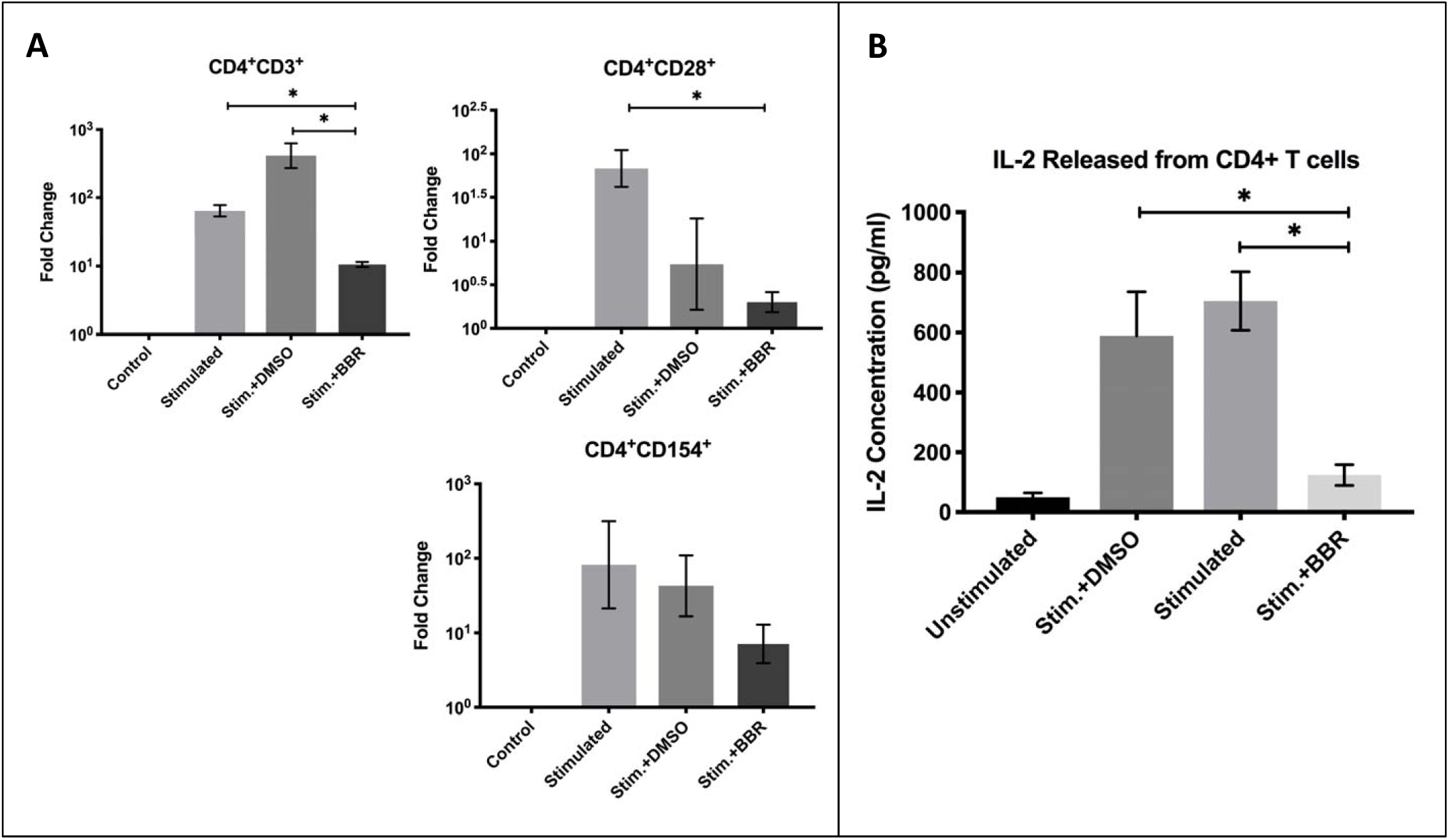
*Berberine suppresses T cell activation* in vitro-Before commencing the CIA animal model, primary T cells from spleen were cultured +/- BBR at 10µM in the presence of anti-CD3 (plate-bound, 5µg/mL overnight before culture) and anti-CD28 (soluble, 2µg/mL) for 5 days. **A.** Populations of CD4+CD3+ T helper cells and expression of co-stimulatory molecules CD154 and CD28 were measured via flow cytometry. Statistical comparisons made with the Kruskal-Wallis test with Dunn’s multiple comparisons (*p<0.05). **B.** Cell culture supernatants were assayed for IL-2, a canonical indicator of T cell activation and autocrine signaling. Statistical comparisons made with the Kruskal-Wallis test with Dunn’s multiple comparisons (*p<0.05).

