## Supplementary figures and images for "Berberine delays onset of collagen induced arthritis through T cell suppression"

### Supplementary Figure 1

#### Slide 1
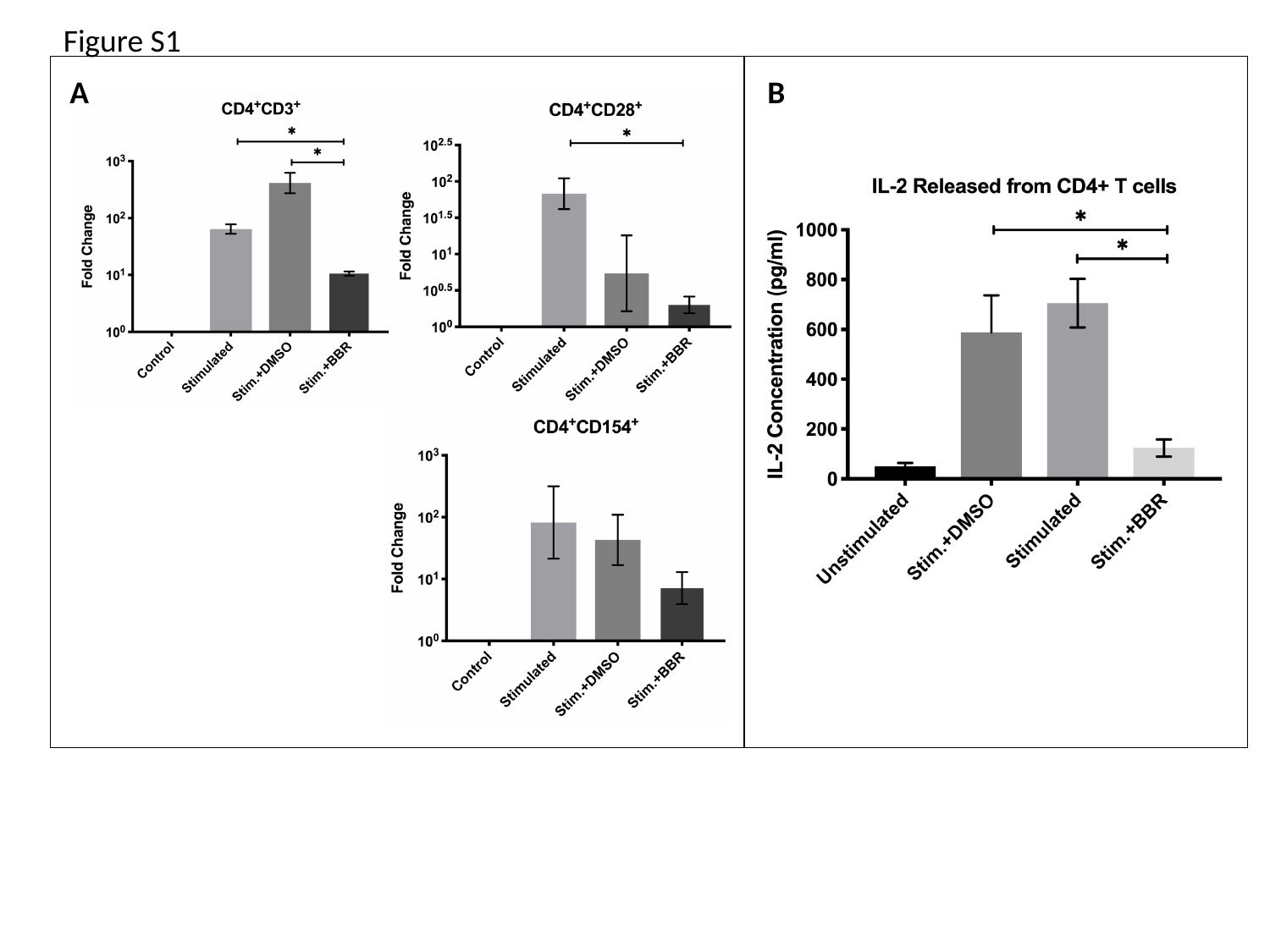

Figure S1
A
B
